# TRPV1-cardiac afferent ablation after completed myocardial infarction prevents arrhythmogenic remodeling

**DOI:** 10.64898/2026.08.31.748451

**Authors:** Kiyoshi Masuyama, Youmei Shen, Wei-Hsin Chung, Abdullah Sarkar, Daisetsu Aoyama, Devaki Abhyankar, Pradeep S. Rajendran, Ronald Challita, Rami Aladham, Lily Defelice, Shumpei Mori, Maureen A. Su, Jeffery L. Ardell, Olujimi A. Ajijola

**Author notes:** **Corresponding author contact information**: Olujimi Ajijola, MD, PhD, UCLA Cardiac Arrhythmia Center Department of Medicine-Cardiology 100 Medical Plaza, Suite 660, Los Angeles, CA 90095. These authors contributed equally to this work (Kiyoshi Masuyama, Youmei Shen).

## Abstract

**Structured Abstract: Background:** Therapies to prevent ventricular arrhythmias after completed myocardial infarction (MI) remain limited. Although TRPV1 afferent ablation at acute MI improves cardiac remodeling, the effect of delayed subacute targeting remains unknown.

**Objective:** We investigated whether ablating cardiac TRPV1 afferents during the subacute post-MI window mitigates structural, electrophysiological, and neuro-cardiac axis remodeling to suppress ventricular arrhythmias.

**Methods:** Yorkshire pigs underwent sham surgery or anterior MI creation. Two weeks post-MI, animals were randomized to percutaneous epicardial resiniferatoxin (RTX, for cardiac-selective TRPV1 afferent depletion) or vehicle administration. Four weeks later, terminal studies assessed the effects of RTX on cardiac structure and function, ventricular arrhythmogenesis, and neuro-cardiac axis remodeling using *in vivo* electrophysiologic mapping, real-time neurotransmitter sensing, immunohistochemistry, and transcriptomic profiling.

**Results:** Cardiac TRPV1 afferent depletion was confirmed by blunted responses to TRPV1 agonists. RTX-treated animals exhibited improved left ventricular function, reduced end-diastolic diameter, and suppressed ventricular tachycardia/fibrillation (VT/VF) inducibility. Endocardial electroanatomic mapping revealed improved VT/VF electrophysiologic correlates in RTX-treated animals, including fewer deceleration zones and late potentials. Epicardial multielectrode mapping demonstrated reduced electrophysiologic heterogeneity in the scar border zone. Real-time release of adrenergic neurotransmitters (noradrenaline and neuropeptide Y) was normalized during sympathoexcitation in RTX-treated animals. Histologically, RTX treatment attenuated scar border zone myocardial fibrosis and sympathetic nerve sprouting, while suppressing T cell infiltration in cardiac sensory ganglia. Bulk RNA-sequencing of stellate ganglia revealed downregulation of adrenergic genes.

**Conclusion:** Cardiac TRPV1 afferent ablation post-completed MI alters disease trajectory by mitigating structural, functional, and neuro-cardiac axis remodeling. Targeting cardiac TRPV1 afferents represents a promising subacute post-MI therapeutic strategy.

**Unstructured Abstract:** Therapies to prevent ventricular arrhythmias following completed myocardial infarction (MI) remain limited, and the impact of targeting cardiac TRPV1 afferents during the subacute phase is unknown. We evaluated whether epicardial resiniferatoxin (RTX) administration two weeks post-MI in swine mitigates structural, electrophysiological, and neuro-cardiac remodeling to suppress ventricular arrhythmogenesis. Four weeks post-treatment, RTX-treated animals exhibited improved left ventricular function, reduced chamber dilation, and suppressed ventricular tachycardia/fibrillation (VT/VF) inducibility compared with controls. Electroanatomic and multielectrode mapping revealed fewer deceleration zones, smaller late potential areas, and reduced repolarization heterogeneity in the scar border zone. Real-time neurotransmitter sensing demonstrated normalized adrenergic release during sympathoexcitation. Histological and transcriptomic profiling showed that RTX attenuated border-zone fibrosis, decreased sympathetic nerve sprouting, suppressed sensory ganglion T-cell infiltration, and downregulated stellate ganglion adrenergic gene expression. Cardiac TRPV1 afferent ablation during the subacute post-MI window alters disease trajectory by mitigating structural and autonomic remodeling, representing a promising disease-modifying therapeutic strategy.

## INTRODUCTION

Ventricular tachycardia and ventricular fibrillation (VT/VF) are the leading causes of sudden cardiac death following myocardial infarction (MI).^1–4^ Post-MI, structural, neurochemical, and functional remodeling of cardiac sympathetic neurons drives sympathetic hyperactivation and ventricular arrhythmogenesis.^5–7^ Although early revascularization and optimal medical therapy including neurohormonal blockade reduce post-MI mortality,^8–10^ these interventions remain insufficient in halting disease progression and preventing sudden cardiac death, particularly after a completed MI.^11,12^

The heart is extensively innervated by an interconnected network of sympathetic, parasympathetic, and sensory neurons that collectively regulate cardiovascular function.^5,13^ Emerging data demonstrate that post-MI autonomic dysfunction is part of a systemic multi-organ signaling cascade, where complex heart-brain neuroimmune circuits perpetuate cardiac damage and adverse remodeling.^14^ Current neuromodulatory therapies have predominantly targeted efferent autonomic pathways by inhibiting sympathetic output or enhancing vagal tone.^15–17^ In contrast, therapeutic approaches aimed at blunting sympathovagal imbalance induced by persistent cardiac afferent signaling, specifically via the cardiac sympathetic afferent reflex (CSAR), remain underexplored.^18,19^ The CSAR is driven in part by activation of transient receptor potential vanilloid 1 (TRPV1) channels expressed on cardiac afferent fibers.^20^ Similar to sympathetic nerve remodeling, sensory fibers display hypersensitivity at the scar border zone, suggesting that cardiac afferent signaling is enhanced following MI. ^21,22,23^ Recent work highlights that targeting these spinal nociceptive afferent pathways can disrupt the chronic autonomic remodeling that ensues post-MI.^24^

Resiniferatoxin (RTX) is an ultra-potent TRPV1 agonist that can selectively deplete TRPV1-positive nociceptive afferent fibers.^25,26^ Prior studies have shown that RTX abolishes sympathetic reflex responses to cardiac stress in the normal heart,^27^ and when administered at the time of MI, reduces VT/VF susceptibility, limits infarct size, and preserves left ventricular (LV) function in a porcine model.^19^ However, whether delayed cardiac afferent ablation can still exert beneficial effects once post-MI pathological remodeling has been initiated, particularly in the setting of completed, non-reperfused MI, remains unknown.

To address this question, we evaluated the efficacy of percutaneous epicardial RTX administration two weeks after completed MI in a porcine model. Animals were randomized in this subacute stage to evaluate the therapeutic potential of delayed TRPV1 afferent depletion on neuro-cardiac remodeling and ventricular arrhythmogenesis. By targeting a clinically relevant subacute window currently unaddressed by existing therapies, this study introduces a novel neuromodulatory strategy to alter the disease trajectory following completed MI.

## METHODS

All animal studies were approved by the UCLA Animal Research Committee (ARC) and conducted in accordance with ethical guidelines.

### Data availability

All data presented in this article are available from the corresponding author upon reasonable request.

### Study design

This was a randomized, *in vivo* porcine study designed to evaluate the effects of cardiac-selective TRPV1 afferent ablation using RTX during the subacute phase of a completed MI. Anterior MI was induced in male and female Yorkshire pigs (Sham MI = 11, MI = 25, ∼45kg) by left anterior descending (LAD) coronary artery occlusion. Two weeks post-MI, animals were randomized to receive epicardial injection of RTX or saline (n=14 and 11, respectively). For all procedures, animal subjects were sedated with Telazol (4-8 mg/kg, intramuscular), intubated, and maintained under general anesthesia with inhaled isoflurane (2%) in supplemental oxygen (2 L/min). Surface electrocardiogram, arterial blood pressure, and body temperature were continuously monitored.

### MI model

An anteroapical MI was created by microbead occlusion of the LAD artery distal to the 2^nd^ diagonal branch, as previously described.^16–18^ Briefly, under fluoroscopy guidance, an AL-0.75 catheter (6Fr, Amplatz-type catheter, Medtronic) was used to advance an angioplasty balloon (2.5mm, Abbott Vascular, Armada 18, Abbott Vascular) over a 0.014” Hi-Torque Balance Middleweight Universal guidewire (Abbott Vascular). The balloon was then inflated at the 2^nd^ diagonal branch of the LAD artery, and 4.0 mL of polystyrene-microspheres (90 μm diameter, Polyscience Inc.) were slowly injected. Following injection and balloon deflation, repeat angiography demonstrating loss of blood flow to the distal LAD, combined with acute surface electrocardiogram ST-segment elevation, confirmed successful MI. If the animal developed VT/VF, external cardioversion/defibrillation and standard cardiopulmonary resuscitation were performed.

### Epicardial RTX administration

Methylprednisolone sodium succinate (0.15 mg/kg, intramuscular) and famotidine (40mg, per oral) were given one day prior to RTX administration. The left sternocostal angle was incised, and percutaneous access to the intrapericardial space was obtained using a Tuohy needle under fluoroscopic guidance. A small amount of contrast agent was injected to ensure the advancement of the needle tip, and intrapericardial access was confirmed by advancing a J-tip guidewire that conformed to the cardiac silhouette under fluoroscopy. A multipurpose catheter was then advanced over the wire to aspirate pericardial fluid and administer 10 mL of RTX. After a 10-minute incubation period, the RTX was aspirated, and methylprednisolone (50 mg) was injected intrapericardially to prevent pericardial adhesions. Post-procedural analgesia with buprenorphine (0.02 mg/kg, intramuscular) was administered starting 6 hours post-procedure and then every 24 hours for 3 days. Famotidine (40 mg/day, per oral) was also continued for 3 days. The same protocol was applied to both the MI + Saline and Sham MI groups, with normal saline administered in place of RTX.

### Terminal experimentation

Terminal studies were performed 4 weeks after the epicardial intervention. After sedation and intubation, transthoracic echocardiography was performed with the chest closed. Arterial blood pressure was monitored continuously and a pressure catheter (SPR350, Millar Inc) was inserted into the LV to monitor LV pressure in real time. Arterial blood gases were evaluated every 90 minutes and ventilation parameters were adjusted or sodium bicarbonate was administered as needed to maintain physiological stability. The thorax and heart were exposed through a median sternotomy. Following completion of surgical procedures, isoflurane was gradually tapered off and switched to α-alose (6.25 mg/125 mL; 1mL/kg for bolus, 20-35mL/kg intravenous or titrated to effect for maintenance). The right stellate ganglia (SG) and both the right and left vagus nerves were exposed for right sympathetic nerve stimulation (RSNS) and bilateral vagal nerve stimulation (BVNS), respectively. At the end of the study, animals were euthanized under deep α-chloralose sedation and induction of ventricular fibrillation.

#### Nerve stimulation

After equilibration from surgical procedures and transition to α-chloralose, bipolar platinum nerve cuffs (Microprobes) were placed around the right SG and the right and left vagus nerves. Nerve stimulation was performed using a Grass S88 stimulator interfaced through a PSIU6 photoelectric constant-current isolation unit (Grass Instruments). For the RSNS, threshold was defined as the current required to elicit a 15% increase in heart rate during a 20-second stimulation train at a frequency of 4 Hz and pulse width of 4 ms. For VNS (right and left), the threshold was defined as the current required to elicit a 15% decrease in heart rate during a 5-second stimulation train at a frequency of 10Hz and pulse width of 1ms. RSNS and BVNS were performed at 1.5 times threshold for 30 seconds.

#### Epicardial electrophysiologic mapping

Epicardial unipolar electrograms were continuously recorded using a 64-electrode array (5.5 mm inter-electrode spacing; MappingLab,) positioned over the LV scar-border zone. Activation recovery interval (ARI), a surrogate of local action potential duration, was calculated as the time difference between repolarization time and activation time (AT) using the EMapScope software (MappingLab). AT was measured from the beginning to the minimal dV/dt of the depolarization wavefront of the unipolar electrogram. Repolarization time was measured from the initial deflection to the maximal dV/dt of the repolarization wavefront. ARIs were corrected for heart rate using Bazett’s method. ARI variability was quantified as the standard deviation of ARI across all 64 channels. ARI dispersion was defined as the difference between the maximum and minimum ARI values. The maximum ARI gradient, reflecting the steepest local change in repolarization duration, was calculated as the largest ARI difference between adjacent electrodes divided by the inter-electrode distance.^28^ These parameters were used to assess spatial heterogeneity in repolarization, which is known to be associated with arrhythmogenic risk.^28^

#### Physiologic responses to TRPV1 agonists

During terminal experiments, MI + Saline and MI + RTX animals were exposed to epicardial bradykinin (40 and 80 mg/mL) and capsaicin (80 mg/mL) to evaluate CSAR. Drugs were applied using an adsorbed patch to the anterior aspect of the ventricles for 1 min. Bradykinin was applied before capsaicin due to sensory peptide depletion by capsaicin. The epicardium was washed with warm saline 10 minutes after each application and a 20-minute waiting period was allowed for recovery of the baseline hemodynamic function.

#### Ventricular arrhythmia inducibility

Arrhythmia susceptibility was tested using both cesium chloride (CsCl) and programmed electrical stimulation (PES). CsCl (84mg/kg) was injected intravenously via the femoral vein, and then flushed with 2mL of warm saline. The occurrence of ventricular arrhythmia was monitored for 10 minutes.

PES was performed using an 8-beat drive train (S1) followed by single (S2) or double (S3) extra stimuli, decremented by 10–20 ms down to the effective refractory period (ERP) or a minimum cycle length (CL) of 200 ms. The S1 drive cycle length (CL) was set to 10% below baseline CL, and initial extra stimulus delivery was set at 75% of S1 CL. If S2 reached ERP without inducing sustained VT/VF, an additional extra stimulus (S3) was introduced. PES was performed using the Micropace (EP320) and Prucka CardioLab System (GE Healthcare). The arrhythmogenicity score was calculated using the following formula: 2*(max PVC beats after S1, S2) + (max PVC beats after S1, S2, S3). If VT/VF was induced during either stimulation, an additional 5 points were assigned. Each stimulation session was scored independently and summed for the final arrhythmogenicity score.

#### Endocardial electroanatomic mapping

During the terminal procedure in sinus rhythm, high-density endocardial electroanatomic mapping was performed in pigs (n = 16) using the Ensite Precision system (Abbott). A multielectrode catheter (Advisor HD Grid, Abbott) was retrogradely advanced into the LV via the aorta. All electroanatomic maps were analyzed offline. The voltage and the local activation time maps were created using the Ensite Precision AutoMap Mapping tool (Abbott).

Bipolar voltage maps were generated using peak-to-peak bipolar voltage signals. Core scar was defined as areas with voltage < 0.5 mV, border zone as 0.5-1.5 mV, and healthy myocardium as > 1.5 mV. Total scar area was calculated as the sum of the core scar and border zone areas.

Isochronal late activation mapping (ILAM) was constructed to visualize the completion of local activation using automated late annotation (EnSite Precision, last deflection, Abbott), as previous described^29^. The thickness of each isochrone reflected local conduction velocity, with wider isochrones indicating faster propagation and narrower, crowded isochrones indicating slower conduction. Deceleration zones were defined as regions with isochronal crowding with > 3 isochrones within a 1-cm radius.^29^ Late potentials were defined as any low-voltage electrograms displaying single or multiple discrete delayed components occurring after the end of the surface QRS complex. All areas on the electroanatomic maps were measured using Ensite system’s surface area and measurement tools.

### RNA sequencing

Following euthanasia, the left SG and dorsal root ganglia (DRG) were rapidly excised, rinsed in saline, and snap-frozen in liquid nitrogen. RNA was isolated using the RNeasy Mini Kit (Qiagen Sciences, Germantown, MD). RNA quality was measured using the Agilent TapeStation Systems (Agilent Technologies Inc, Santa Clara, CA), while RNA concentration was measured with Nanodrop spectrophotometer (ThermoFisher Scientific). Libraries were prepared for RNA sequencing with KAPA mRNA HyperPrep Kit (Roche). The workflow consisted of mRNA enrichment, cDNA generation, end repair to generate blunt ends, A-tailing, adaptor ligation, and PCR amplification. Different adapters were used for multiplexing samples in one lane. Sequencing was performed on Illumina NovaSeq X Plus 10B for 50-bp single-read run. Data was analyzed with Illumina software. Functional enrichment analysis of the gene profile (GO biological process and KEGG pathway analysis) was performed using the Database for Annotation, Visualization and Integrated Discovery (DAVID) 2021.

### Statistical analyses

Data are presented as mean ± standard error of the mean (SEM) for normally distributed continuous variables, or as median and interquartile range (IQR) for non-normally distributed or ordinal variables. Categorical variables were expressed as counts and percentages. Normality of continuous variables was assessed using the Shapiro-Wilk test. For comparisons between two groups, normally distributed variables were analyzed using the unpaired Student’s t-test, while non-normally distributed or ordinal variables were compared using the Mann–Whitney U test. For comparisons among more than two groups, one-way ANOVA was used for normally distributed variables with equal variances (assessed by the Brown-Forsythe test), and Welch’s ANOVA was applied if variances were unequal. For non-normally distributed variables, the Kruskal–Wallis test was used. Multiple comparisons were controlled by using Tukey’s or Dunnett’s test, as appropriate. Categorical variables were compared using the Chi-square test or Fisher’s exact test, as appropriate. Ventricular arrhythmias induced by CsCl infusion were analyzed using two-way repeated measures ANOVA in GraphPad Prism. Time (1–4 minutes post injection) and treatment group (MI + Saline vs MI + RTX) were treated as fixed factors, with subject as the repeated measure. The Geisser–Greenhouse correction was applied to account for violations of sphericity. The interaction between time and group was included in the model. A two-sided p-value of <0.05 was considered statistically significant. Statistical analysis and preparation of graphs were conducted using GraphPad Prism (GraphPad Software Inc., La Jolla, CA, USA).

## RESULTS

### Epicardial RTX application induces degeneration of TRPV1 afferent fibers

Cardiac-selective TRPV1-expressing fibers were targeted using a percutaneous epicardial approach to transiently deliver RTX or vehicle to the epicardium following completed MI. For a clinically relevant subacute window, RTX was applied 2 weeks post-MI (Figure 1A). A dose of 5.0 μg/ml was selected based on preliminary dose ranging studies (evaluating 2.5, 5.0, and 12.5 μg/ml) (Supplemental Figure 1 A-F). During percutaneous epicardial application of RTX, animals consistently exhibited brief vagal responses, characterized by transient reductions in heart rate and blood pressure within 1–2 minutes of application, confirming drug effect (Supplemental Figure 2A and 2B). Four weeks later, challenge with high-dose TRPV1 agonists bradykinin (40 μg/ml, 80 μg/ml) and capsaicin (80 μg/ml) confirmed that TRPV1 afferent-mediated sympathetic reflex responses were functionally abolished (Figure 1B and 1C). We confirmed that the effects of RTX treatment were limited to afferent nerve depletion, as there was no impact on efferent sympathetic or parasympathetic responses (Supplemental Figure 3A-3C). We also examined the distribution of sensory neurons in the DRG to investigate whether afferent depletion led to loss of DRG neurons. We did not identify any regions of cell loss within DRG, and further, percentage of CGRP-positive cells (key mediators or cardiac afferent neurotransmission) did not change following TRPV1 afferent ablation (60.21 ± 5.11 vs 56.45 ± 11.28%; p = 0.66) (Supplemental Figure 3D and 3E).

**Figure 1.**
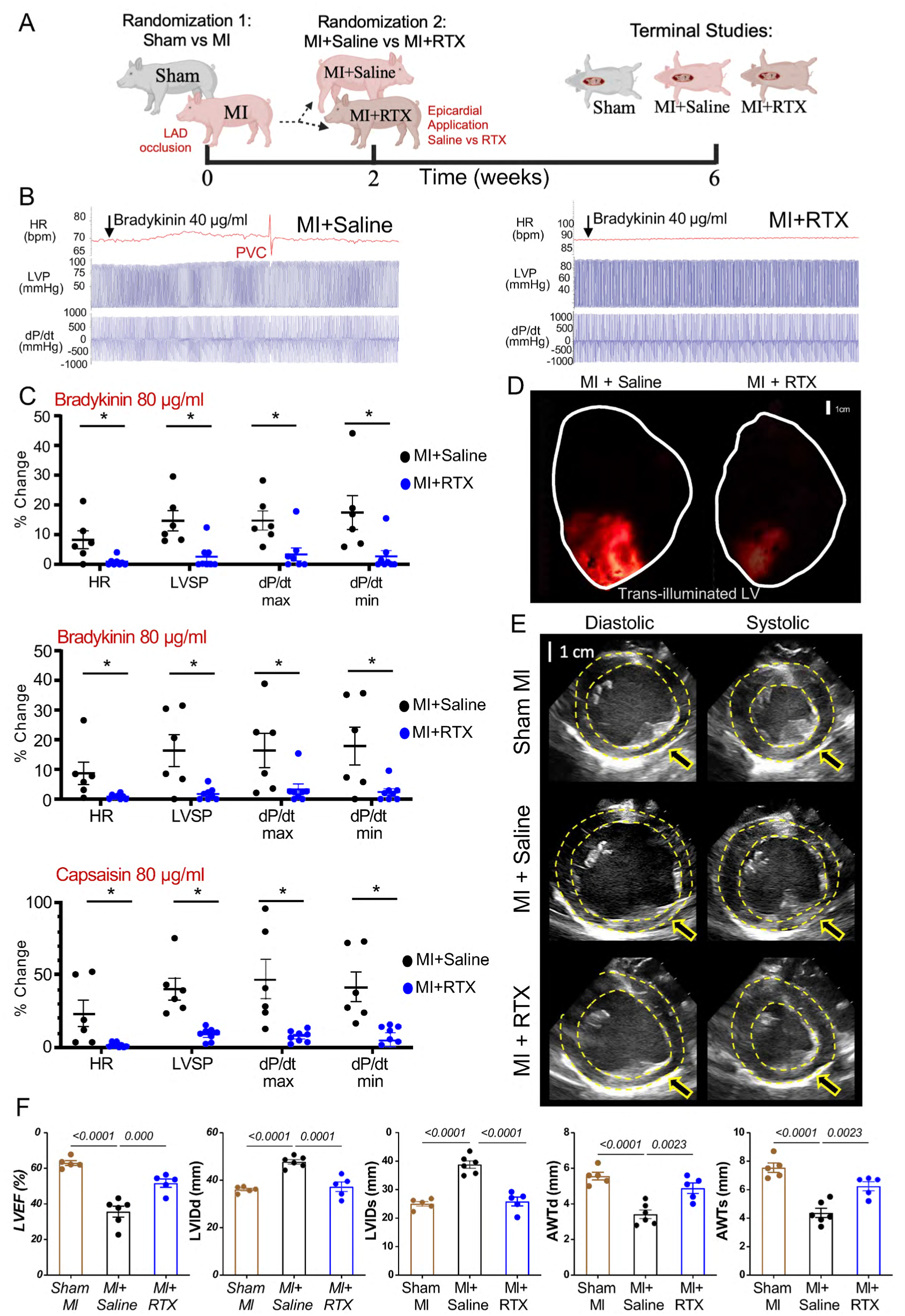
Cardiac afferent TRPV1 depletion by resiniferatoxin attenuates post-myocardial infarction left ventricular dysfunction. (A) Schematic image of the experimental protocol. (B) Representative images of the hemodynamic reaction to application of TRPV1 agonist to both groups. (C) Epicardial TRPV1 afferent depletion abolished sympathetic reaction under bradykinin 40μg/ml, 80μg/ml, and capsaicin 80μg/ml. MI + Saline animals showed hemodynamic sympathetic response, but MI + RTX animals did not display this response. n = 6-9/group. Analyzed by Mann-Whitney U-test. (D) Representative heart images from MI + Saline and MI + RTX subjects. Light highlights the scar area of the left ventricle. RTX treatment reduced scar area (scale bar = 1 cm). (E) Representative left ventricular short axis diastole and systole echocardiographic images from three groups (scale bar = 1 cm). (F) Quantified data among Sham MI, MI + Saline, and MI + RTX animals. The MI + Saline group exhibited significant impairment in left ventricular ejection fraction (LVEF), dilation of the left ventricular internal diameter at end-diastole (LVIDd) and end-systole (LVIDs), and anterior wall thinning in both diastole (AWTd) and systole (AWTs) compared to Sham MI animals. RTX treatment markedly attenuated these structural and functional abnormalities post-MI, improving LVEF, reducing chamber dilation, and preserving anterior wall thickness. Data are presented as mean ± SEM and analyzed by one-way ANOVA followed by Dunnett’s multiple comparisons test. N = 5-6 per group. *P<0.05; **P<0.01; ***P<0.001.

### Epicardial RTX application after completed MI mitigates cardiac structural abnormalities and rescues cardiac contractile function

Visually, epicardial scar area appeared smaller in the MI + RTX animals compared to vehicle-treated animals (Figure 1D). Consistent with this, echocardiography revealed that animals in the MI + saline group had significantly impaired LV function and structure compared to the sham MI group, including a lower LV ejection fraction (LVEF; 35.6 ± 3.1% vs 63.0 ± 1.4%, p < 0.001) and larger LV internal diameter at end-diastole (LVIDd; 47.7 ± 0.9 vs 36.0 ± 0.6 mm, p < 0.001), and at end-systole (LVIDs; 38.8 ± 1.3 mm vs 25.0 ± 0.8 mm, p < 0.001) (Figure 1E and 1F). The LV anterior wall thickness was also significantly reduced in both diastole (AWTd; 3.4 ± 0.2 mm vs 5.6 ± 0.2 mm, p <0.001) and systole (AWTs; 4.4 ± 0.3 mm vs 7.5 ± 0.3 mm, p <0.001). In contrast, the MI + RTX group showed attenuated adverse remodeling following completed MI, with improved LVEF (51.7 ± 2.3% vs 35.6 ± 3.1%, P = 0.001), smaller LVIDd (36.0 ± 0.6 mm vs 47.7 ± 0.9 mm, P < 0.001) and LVIDs (25.0 ± 0.8 mm vs 38.8 ± 1.3 mm, P < 0.001), and greater AWTd (4.9 ± 0.3 mm vs 3.4 ± 0.2 mm, P = 0.002) and AWTs (6.2 ± 0.3 mm vs 4.4 ± 0.3 mm, P = 0.002) (Figure 1E and 1F). These findings demonstrate that RTX application after completed MI improved cardiac structure and function.

### Cardiac TRPV1 afferent depletion reduces VAs after MI

To evaluate whether the structural benefits derived from depletion of cardiac TRPV1 afferents reduces VAs after completed MI, we probed ventricular arrhythmogenesis using complementary approaches. First, CsCl, a nonspecific potassium channel antagonist that transiently elicits afterdepolarization-induced triggered activity, was administered. In MI + saline animals, a higher frequency of premature ventricular contractions (PVCs) and non-sustained ventricular tachycardia (NSVT) episodes were observed, especially in the first 4 minutes post-injection (Figure 2A). In contrast, in the MI + RTX group, fewer PVCs and NSVT episodes were observed (Figure 2B and 2D). The number of PVCs at 1, 2, 3, and 4 minutes after injection was significantly reduced in the MI + RTX group compared to the MI + saline group (61.0 ± 16.3 vs 20.4 ± 3.6 beats at 1 min post injection, 46.0 ± 11.5 vs 7.8 ± 5.1 beats at 2 min, 32.6 ± 8.5 vs 2.9 ± 1.5 beats at 3 min, 22.2 ± 10.0 vs 2.4 ± 2.0 beats at 4 min; P < 0.001). The total number of PVCs and NSVT episodes trended to be lower in the MI + RTX group compared to the MI + Saline group (PVCs: 34.1 ± 9.0 vs 169.6 ± 45.6 beats, p = 0.06; NSVT episodes: 2.8 ± 0.7 vs 15.6 ± 3.9, p = 0.07) (Figure 2C-2E).

**Figure 2.**
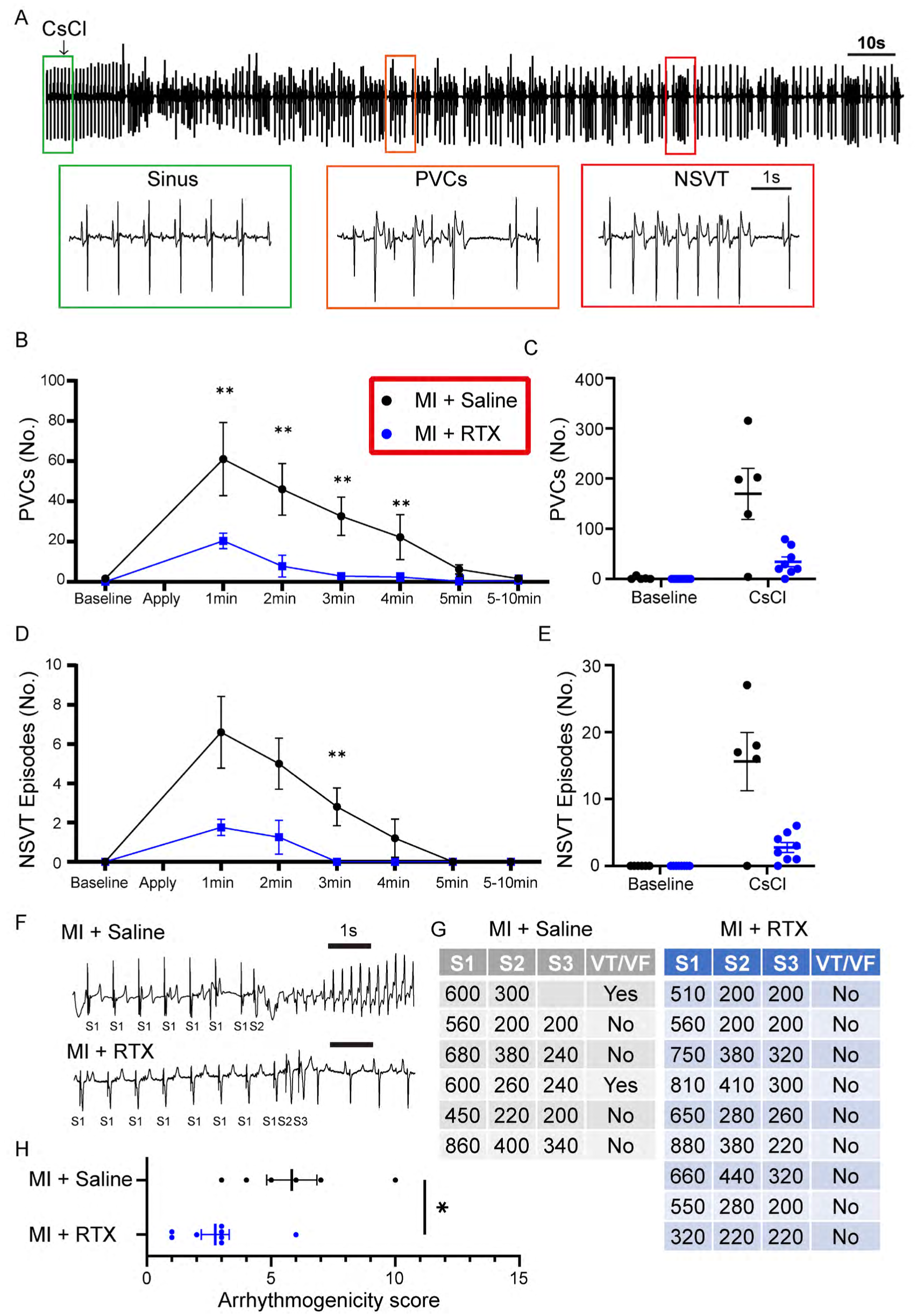
Cardiac TRPV1 afferent depletion reduces ventricular arrhythmogenesis after completed myocardial infarction. (A) Representative arrhythmic change after cesium chloride (CsCl) intravenous administration 84 mg/kg) (scale bar = 10s). Green, orange, and red boxes are zoomed in sinus rhythm, frequent premature ventricular contractions (PVCs), and non-sustained ventricular tachycardia (NSVT), respectively (scale bar = 1 s). (B, D) Quantified data of the number of PVCs and NSVT episodes time dependently. (C, E) Comparison of total number of PVCs and NSVT episodes at the baseline and 10 mins after CsCl injection. (F) Representative tracing of ventricular tachycardia (VT) induced by programmed electrical stimulation in MI + Saline with 1 extra stimuli (S2) but not in MI + RTX group. (G) Across subjects, VT induction was reduced in MI + RTX subjects compared with MI + Saline. (H) Arrhythmogenicity index, calculated from the number of PVCs after extra stimuli was reduced in MI + RTX subjects vs MI + Saline. (n= 6,9 respectively). *P<0.05; **P<0.01; ***P<0.001 by robust repeated measure analysis of variance model and Mann-Whitney U test.

Ventricular arrhythmogenesis was further assessed by programmed extra-stimulation (PES) from a right ventricular endocardial catheter. VT/VF was induced in 2 of the 6 MI + Saline subjects but in 0 of the 9 MI + RTX subjects (Figure 2F and 2G). In addition, the arrhythmogenicity score (comprised of the ease of arrhythmia induction, PVCs, and brief episodes of VT) was significantly higher in the MI + Saline group compared to the MI + RTX group (5.8 ± 0.9 vs 2.8 ± 0.5, p = 0.01) (Figure 2H). Together, these findings indicated that depleting cardiac TRPV1 afferents following completed MI suppresses ventricular arrhythmogenesis.

### Anti-arrhythmic actions of TRPV1 afferent depletion are mediated by reducing scar border zone size and electrophysiologic heterogeneity

Next, we sought to identify the antiarrhythmic mechanisms of cardiac afferent depletion following completed MI. To identify clinically relevant mechanisms, we performed high-density endocardial electroanatomic mapping in a subset of study animals (Sham MI = 5, MI + Saline = 6, and MI + RTX = 5). Representative bipolar voltage maps, ILAM maps, and electrograms from the MI + saline and MI + RTX groups are shown in Figure 3A. No low-voltage areas were observed in the sham MI animals. However, compared to MI + Saline animals, RTX-treated animals exhibited significantly smaller total scar (13.4 ± 2.1 mm² vs 26.3 ± 4.8 mm², p = 0.048) and border zone areas (8.4 ± 1.5 mm² vs 16.9 ± 3.1 mm², p = 0.047), with no significant difference in core scar areas (5.4 ± 1.4 mm² vs 9.4 ± 2.0 mm², p = 0.149). This suggests that primary actions of TRPV1-afferent depletion after MI is to limit border zone expansion.

**Figure 3.**
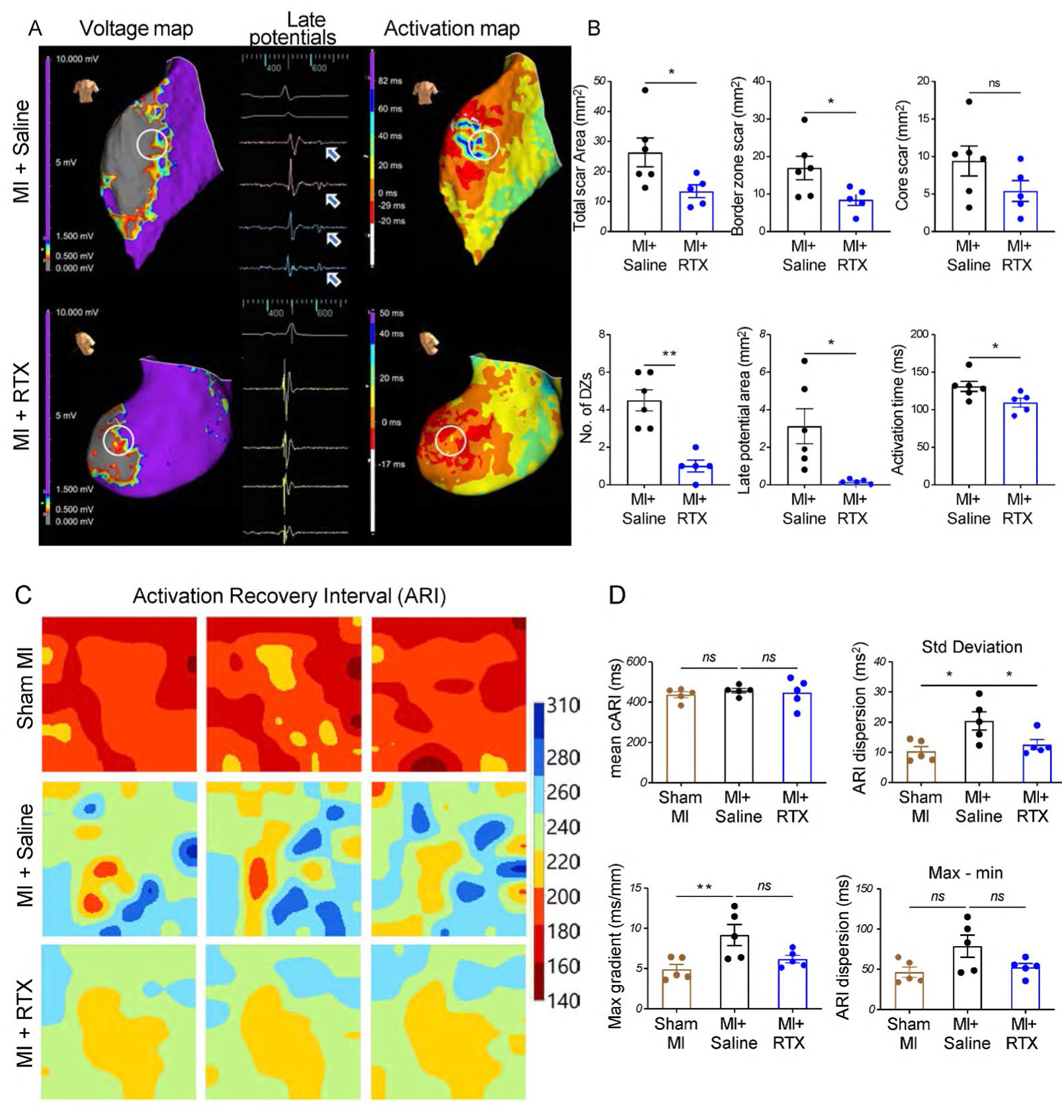
Anti-arrhythmic actions of TRPV1 afferent depletion are mediated by reducing scar border zone size and electrophysiologic heterogeneity. (A) Representative voltage maps, isochronal late activation maps (ILAM), and electrograms from MI + Saline and MI + RTX groups. In voltage maps, areas with bipolar voltage >1.5 mV (purple) represent healthy myocardium, <0.5 mV (gray) indicate core scar, and 0.5–1.5 mV (color scale) define the infarct border zone. ILAM demonstrated crowded isochrones in the MI + Saline group, reflecting conduction slowing, which was absent in RTX-treated pigs. (B) Quantitative comparison of electrophysiological parameters derived from high-density endocardial mapping (MI + Saline: n=6, MI + RTX: n=5). Voltage maps showed that compared with MI + Saline pigs, RTX-treated animals exhibited significantly smaller total scar area and border zone area, with no difference in core scar area. ILAM showed that the MI + RTX group had fewer deceleration zones, reduced late potential area, and shorter activation time compared with MI + Saline animals. (C) Examples of border zone maps obtained from 1 animal from each group showing activation recovery intervals (ARIs) from these regions. ARI mapping was performed in 15 pigs (n=5 per group) to assess repolarization heterogeneity. (D) There was no significant difference in mean corrected ARI among the groups. However, ARI variability was significantly increased in the MI + Saline group compared with Sham MI and was attenuated by RTX. A similar trend was observed in ARI dispersion. Maximal ARI gradient was significantly elevated in the MI + Saline group compared with Sham MI, and showed a downward trend with RTX treatment. Data are shown as mean ± SEM or median [IQR] as appropriate. Statistical analysis was performed using one-way ANOVA followed by Dunnett’s multiple comparisons test. *P<0.05; **P<0.01; ***P<0.001.

Consistent with this, mapping ventricular activation using the ILAM technique, MI + RTX animals demonstrated significantly fewer deceleration zones (1.0 [0.5, 1.5] vs 4.5 [3.0, 6.0], p = 0.002), reduced LP areas (0.1 ± 0.1 mm² vs 3.1 ± 0.9 mm², p = 0.0178), and shorter activation times indexed to a reference 131 ± 6.6ms vs. 109 ± 5.9ms, p=0.048) (Figure 3B), compared to the MI + saline group. Importantly, these electrophysiologic correlates of arrhythmogenic tissue map to the scar-border zone.

Next, we performed high-resolution epicardial mapping using high-density multielectrode arrays in a subset of study animals (Sham MI: n=5, MI + Saline: n=5, MI + RTX: n=5) to assess ventricular repolarization parameters in the infarct border zone (Figure 3C). At baseline, the mean corrected ARI did not significantly differ among the three groups (437.0 ± 14.6 ms vs 457.1 ± 10.9 ms vs 447.2 ± 31.2 ms for Sham MI, MI + Saline, and MI + RTX, respectively). Similarly, mean activation time (AT), spatial AT dispersion (i.e., gradient), and AT heterogeneity (max-min) were not different across the three groups (Supplemental Figure 4A-4D).

In contrast, ARI instability (as measured by beat-to-beat variability) was significantly increased in the MI + Saline group compared with Sham MI (10.4 ± 1.6 ms vs 20.4 ± 3.0 ms, p = 0.012) and was attenuated by RTX treatment (MI + Saline vs MI + RTX: 20.4 ± 3.0 ms vs 12.5 ± 1.7 ms, p = 0.045). A similar trend was observed in ARI dispersion (Figure 3D). The maximal repolarization gradient, reflecting steepness of spatial repolarization, was significantly elevated in the MI + Saline animals compared to Sham MI (4.9 ± 0.6 ms/mm vs 9.2 ± 1.3 ms/mm, p < 0.001), and MI + RTX animals were not significantly different from sham MI animals (Figure 3D).

### Neuro-cardiac axial mapping reveals restored adrenergic reserve following cardiac-selective afferent depletion

Next, we mapped the neuro-cardiac axis by performing simultaneous cardiac electrical mapping using a 64-electrode array with capacitive immunosensing (CI) and fast-scanning cyclic voltammetry (FSCV) to measure real time release of neuropeptide Y (NPY) and norepinephrine (NE) in the beating heart in vivo (Figure 4A) across scar, border zone, and remote myocardium. In Sham MI, MI + Saline, and MI + RTX animals, we utilized direct electrical stimulation of the right stellate ganglion to induce cardiac sympathoexcitation while measuring NE/NPY release and mapping cardiac electrical function. Consistent with the electroanatomic mapping data we earlier observed, we observed increased ARI dispersion and steepened spatial repolarization gradients in MI + Saline vs. Sham MI animals. These markers of enhanced electrical instability were not present in MI + RTX animals (Figure 4B). Interestingly, we found that selective cardiac TRPV1 afferent depletion altered NPY but not NE release (Figure 4C and 4D) during sympathoexcitation induced by stellate ganglion stimulation. Specifically, while NE release was not different between groups, evoked NPY release appeared reduced in MI + Saline animals compared to sham. This apparent reduction in adrenergic reserve was restored in MI + RTX animals (Figure 4D).

**Figure 4.**
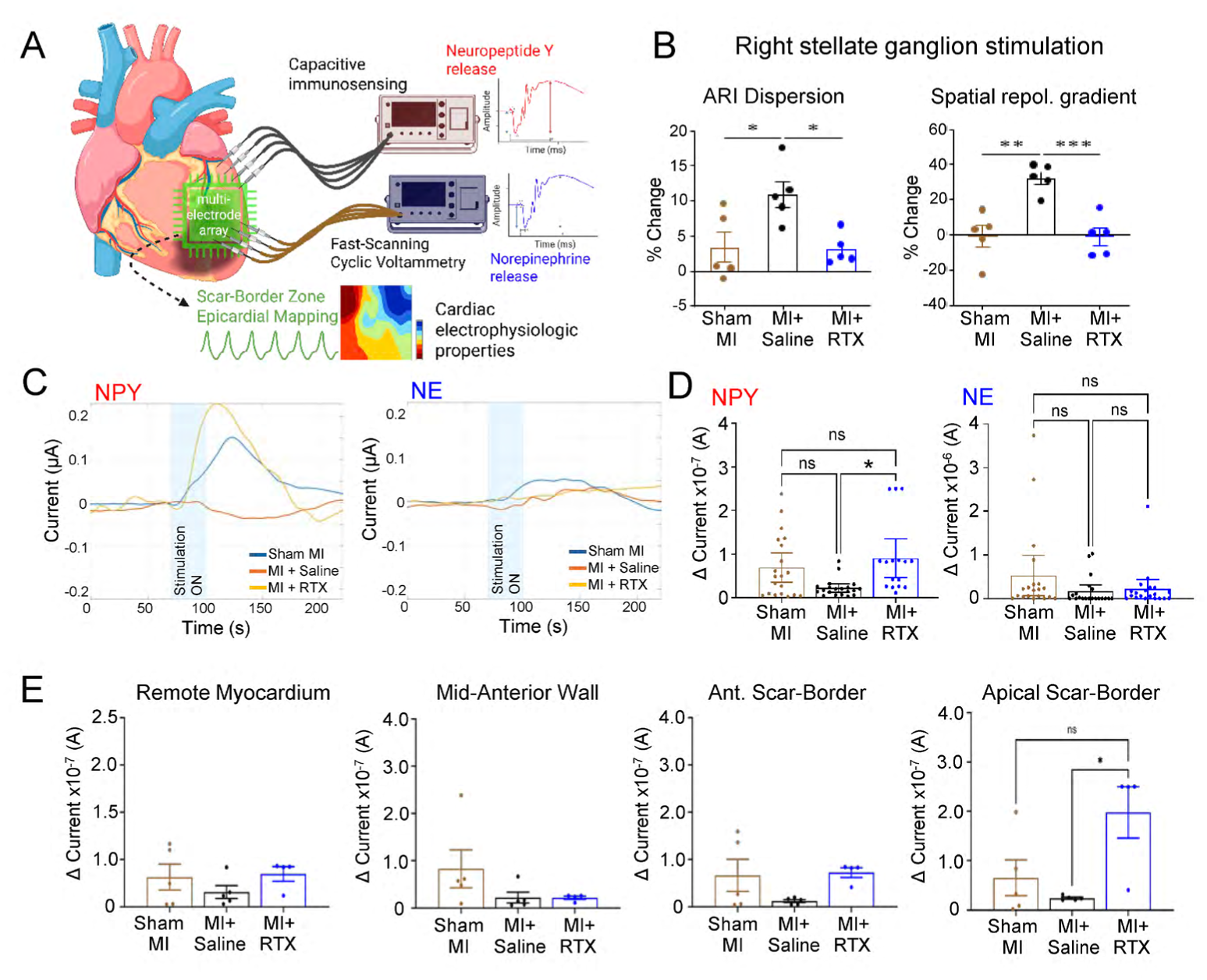
Neuro-cardiac axial mapping reveals restored adrenergic reserve following cardiac-selective afferent depletion. (A) Schematic illustration of the experimental set up for neuro-cardiac mapping. (B) Electrophysiologic heterogeneity (ARI dispersion – indicated by the max – min ARI value across the array, left panel) and spatial repolarization gradients (ms/mm across the array, right panel) in Sham MI, MI + Saline and MI + RTX, n=5/groups). (C) Representative tracings of evoked neuropeptide Y (NPY) and norepinephrine (NE) release during right stellate ganglion stimulation in remote myocardium. (D) Quantified release (i.e. change in current) of NPY and NE at all sites in animals from the three study groups, NPY release is greater in MI + RTX animals compared to MI + Saline, and similar to release levels in Sham MI controls. (E) Quantified release of NPY was greater in the scar border zone but not in other myocardial regions of MI + RTX animals compared MI + Saline. Statistical analysis was performed using one-way ANOVA followed by Dunnett’s multiple comparisons test. *P<0.05; **P<0.01; ***P<0.001.

We then examined regional NPY release patterns across scar-border zone and remote myocardium. As shown in Figure 4E, we found that the return of adrenergic reserve indicated by restored NPY release to or above levels seen in Sham MI animals was localized to the scar-border zones not tissues removed from the infarct site. No spatial differences were seen in NE release patterns across groups (Supplemental Figure 4E). Taken together, these findings indicate that persistent NPY (not NE) release at the scar-border zone is associated with pro-arrhythmic structural remodeling following completed MI and highlight the regional complexity of neural remodeling in the chronically infarcted heart.

### Scar border zone microstructural disarray, fibrosis, and sympathetic nerve sprouting are attenuated by cardiac TRPV1 afferent ablation following completed MI

To examine whether the improvement in scar-border zone electrical abnormalities and restored adrenergic reserve induced by afferent depletion after completed MI were associated with structural correlates, we examined fibrosis and nerve sprouting, known drivers of ventricular arrhythmogenesis, in the scar border zone.

Consistent with bipolar voltage mapping data (Figure 3B), Masson’s Trichrome staining revealed similar dense core scar size in MI + RTX compared with the MI + Saline animals (17.99 ± 2.80% vs 22.29 ± 1.90%; p = 0.10) (Figure 5A and 5B). However, when we examined the scar-border zone in detail, we identified severe microstructural abnormalities (including irregularly distributed and fragmented fibrosis, myocardial fiber disarray and cell density, and surviving cardiomyocytes bordered by scar) which we quantified by a disarray score. Compared to MI + Saline animals, the border zone disarray score was significantly lower in MI + RTX animals (2.2 ± 0.2 vs 1.2 ± 0.2; p = 0.01) (Figure 5C and 5D). In addition, ventricular nerve sprouting, which has been specifically linked to ventricular arrhythmias and sudden cardiac death, was significantly reduced in the MI + RTX group compared with the MI + Saline group, as indicated by the GAP-43-positive area in the border zone normalized to the remote zone (0.6 ± 0.1 vs 1.9 ± 0.3; p = 0.001) (Figure 5E and 5F).

**Figure 5.**
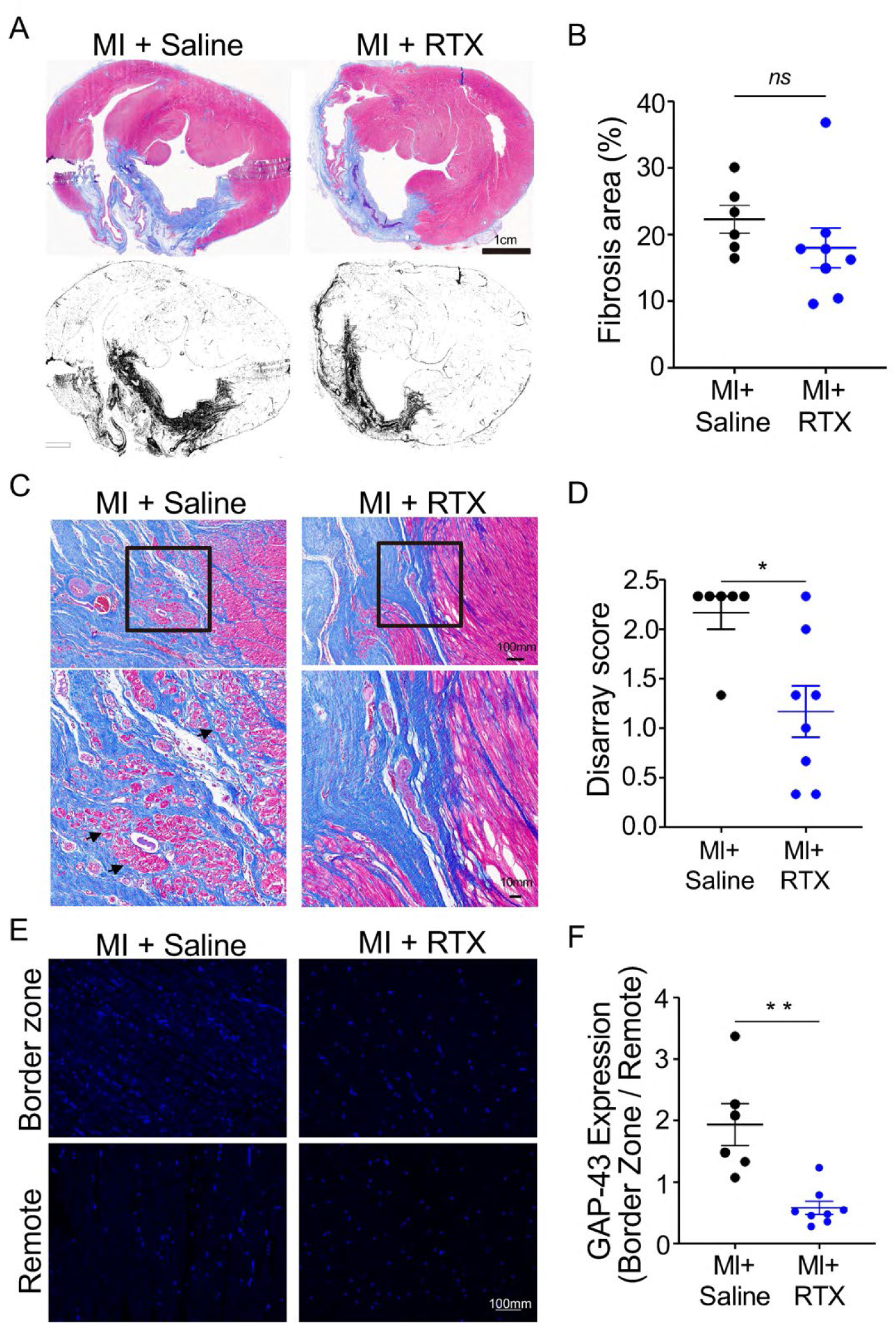
Scar border zone microstructural disarray, fibrosis, and sympathetic nerve sprouting are attenuated by cardiac TRPV1 afferent. (A, B) Representative Masson’s Trichrome image of ventricle showed smaller fibrotic area in MI + RTX group subjects compared to MI + Saline subjects (scale bar = 1cm). (C, D) Representative images of the border zone. Bottom images are magnified images from the box area in the top images (scale bar = 100μm, 10μm, respectively). MI + Saline animals more commonly had myocytolysis, hypertrophy, and decreased muscle band staining, which was less common in MI + RTX animals. (E, F) Representative images of immunoreactivity to growth associated with protein-43 (GAP-43) are shown in the LV border zone and remote zone of MI + Saline and MI + RTX subjects. The normalized GAP-43 calculated by the border zone / remote zone, decreased in the MI + RTX group. n = 6, 8 respectively. *P < 0.05; **P < 0.01 by Mann-Whitney U-test.

### Sensory but not sympathetic neuroinflammation is suppressed by afferent nerve depletion after completed MI

Since VT/VF has been linked to remodeling of the neuro-cardiac axis after MI, we explored whether, following completed MI, selective cardiac afferent depletion attenuates sensory (DRG) and sympathetic neural remodeling. To explore how RTX treatment impacted sensory neurotransmission, we investigated neuroinflammation, a known driver of dysregulated sensory neurotransmission. DRGs were stained with CD3, a marker of T cells (Figure 6A). T cell infiltration in DRGs, measured at intermediate (100X) and high magnification (160X) respectively, was suppressed in the MI + RTX group compared with the MI + Saline group (72.4 ± 8.6 vs. 106.3 ± 7.95, p = 0.02; 32.8 ± 3.2 vs 43.3 ± 1.7; p = 0.03, respectively for 100X and 160x) (Figure 6B and 6C). We also investigated T cell infiltration in sympathetic (stellate ganglia), and observed no differences between MI + Saline and MI + RTX animals 104.7 ± 7.8 vs. 106.2 ± 11.3; p > 0.99) (Supplemental Figure 5A and 5B). Taken together, these data suggest that sensory but not efferent sympathetic neuroinflammation was a target of afferent depletion after MI.

**Figure 6.**
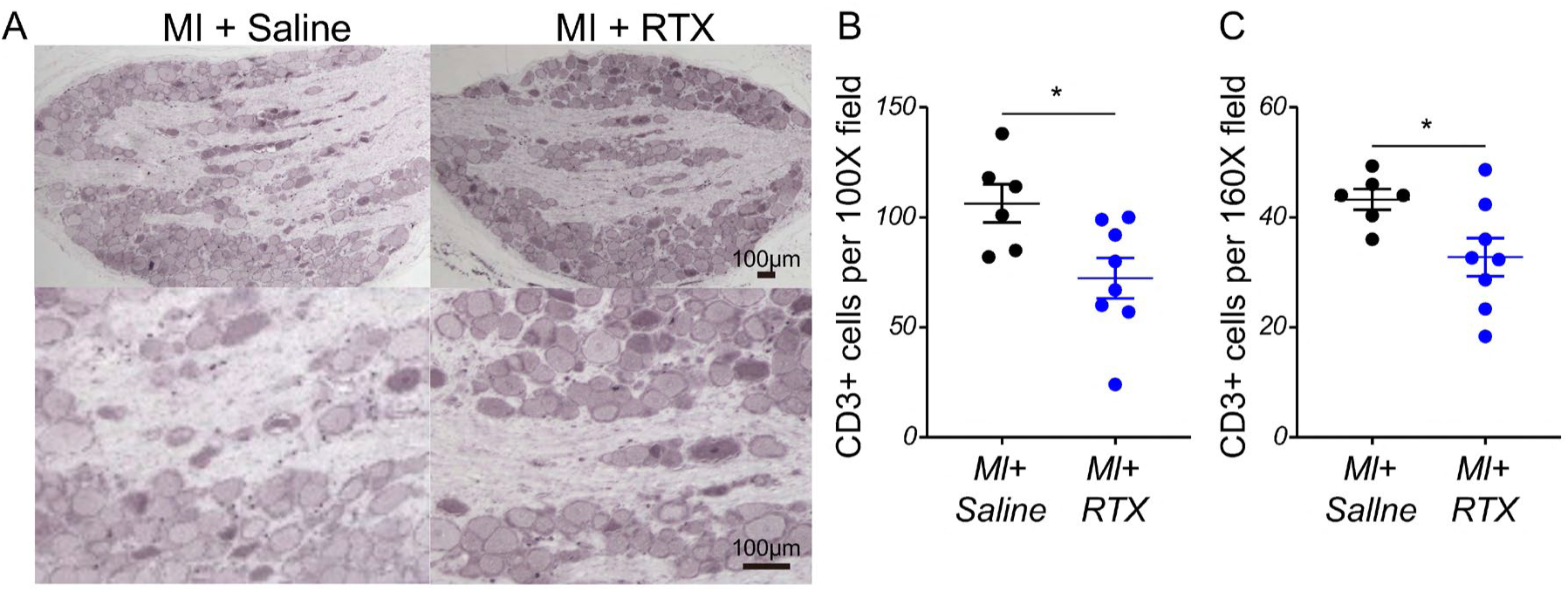
Cardiac afferent TRPV1 depletion mitigates T cell infiltration to DRG. (A-C) Representative images of T cells in DRG using Low-Power Field (LPF) at x 100 magnification and High-Power Field (HPF) at x 160 magnification showed fewer number of positive cells in the MI + RTX group compared with the MI + Saline group (scale bar = 100 μm). (D, E) Representative images of immunoactivity to protein gene product 9.5 (PGP 9.5) and calcitonin gene-related peptide (CGRP) are shown in DRG. No difference was evident in CGRP expression between those groups. n = 6, 8 respectively. *P < 0.05; **P < 0.01 by Mann-Whitney U-test.

### Cardiac TRPV1 afferent depletion suppresses expression of molecular mediators of sympathoexcitation

To clarify how cardiac TRPV1 afferent depletion modulates CSAR, bulk RNA sequencing was performed on SG and DRG tissues from MI + Saline and MI + RTX animals (n = 3/group). In the SG, TH (the rate limiting enzyme in the production of norepinephrine), dopa decarboxylase, and neuropeptide Y (NPY), a proarrhythmic sympathetic co-transmitter, were downregulated in the MI + RTX group compared with the MI + Saline group (Figure 7A, 7B, and 7D). Consistent with this finding, when we examined TH and NPY using immunohistochemistry, we observed trends towards decreased TH and NPY immunoreactivity in MI + RTX compared to MI + Saline animals (Supplemental Figure 5C-5F). Further, we observed that genes involved in axon guidance or long-term synaptic potentiation and extracellular matrix organization were upregulated in the MI + RTX group (Figure 7C). These results suggest that epicardial RTX injection suppresses mediators of adrenergic neurotransmission, including nerve sprouting following completed MI. In the DRG, several genes related to immune response and senescence or p53 pathway were downregulated in MI + RTX group, consistent with our earlier finding that DRG neuroinflammation following completed MI was suppressed by RTX epicardial application.

**Figure 7.**
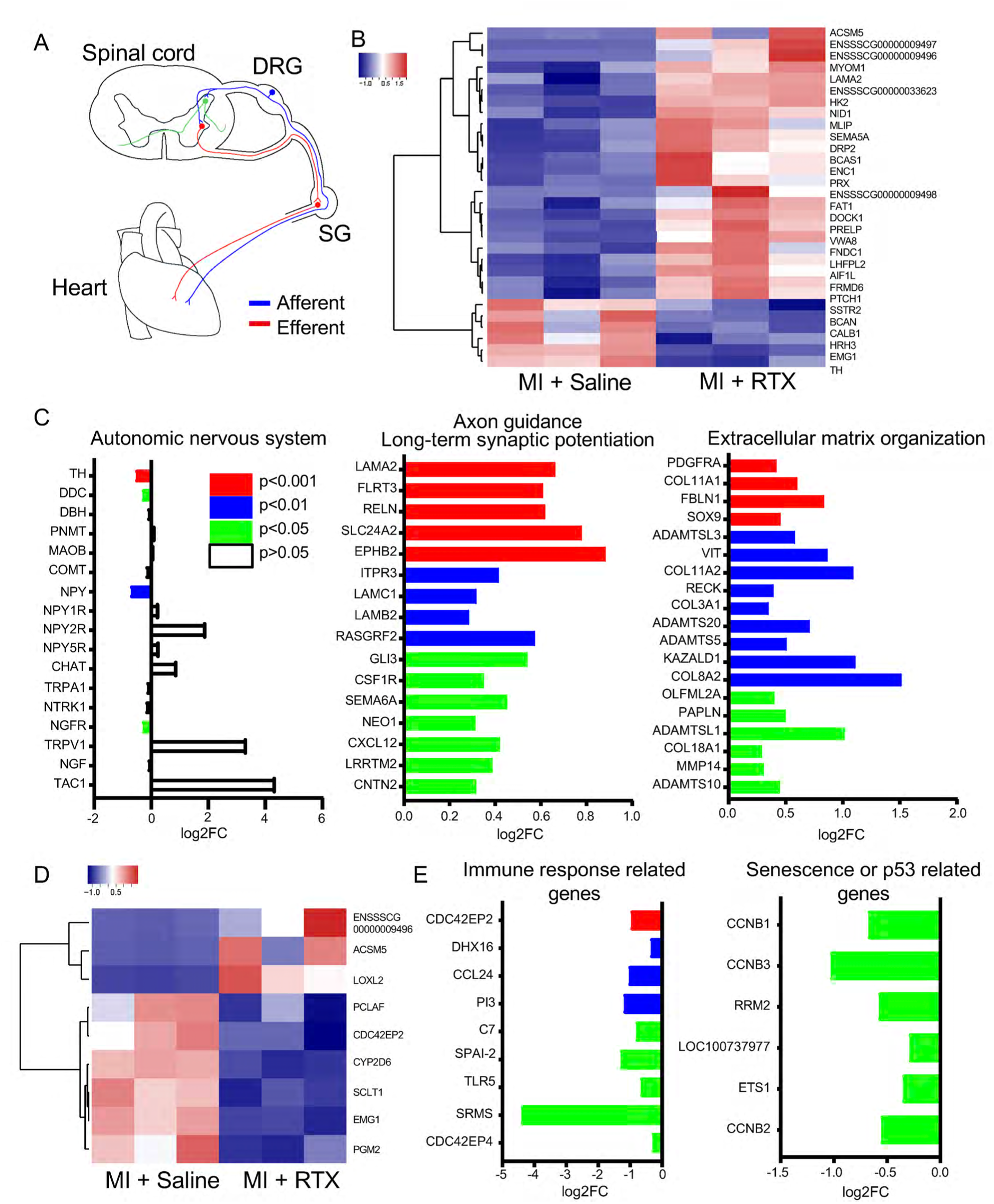
Cardiac TRPV1 afferent depletion suppresses expression of molecular mediators of sympathoexcitation. (A) Schematic image of the cardiac sympathetic afferent reflex loop. (B) Heatmap of significant differentiated genes in SG between MI + Saline and MI + RTX subjects. (C) Some genes involved in autonomic nervous system were downregulated and several genes involved in axon guidance/long-term synaptic potentiation and extracellular matrix organization were upregulated in SG of MI + RTX group. (D) Heatmap of significantly differentiated genes in DRG between MI + Saline and MI + RTX subjects. (E) Several genes involved in immune response and senescence/p53 related genes were downregulated in DRG of MI + RTX group.

**Figure 8.**
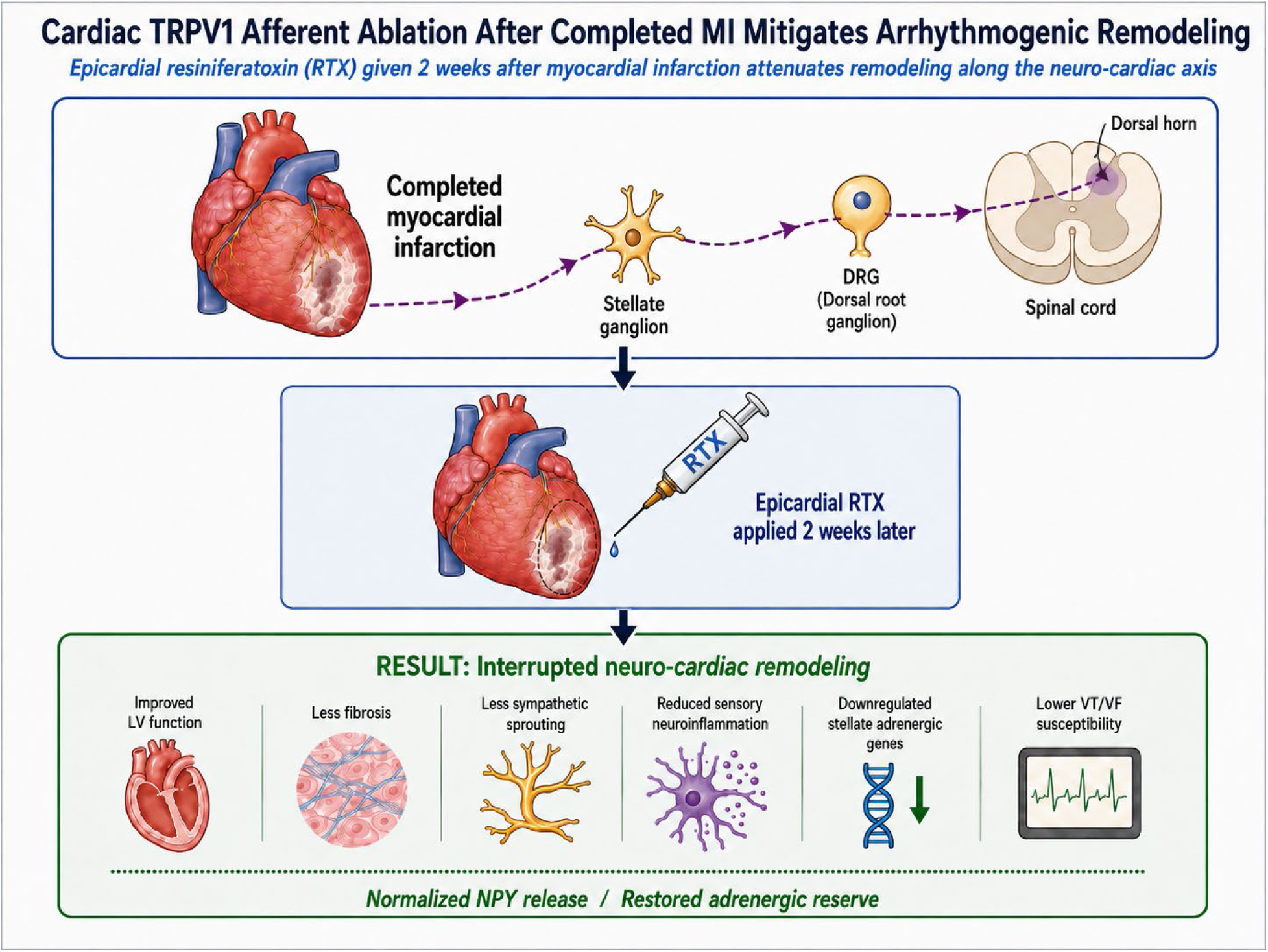
Central illustration. Epicardial RTX administered in subacute post-MI window attenuate neuro-cardiac axis remodeling to suppress ventricular arrhythmogenesis. Visual Abstract

## DISCUSSION

### Major findings

Our study demonstrates that cardiac-selective afferent depletion with epicardial RTX, administered two weeks after a completed MI, attenuates adverse cardiac remodeling, reduces VT/VF, and limits neuro-cardiac axis plasticity. Importantly, we show for the first time that delayed RTX treatment restores regional adrenergic reserve in the infarct border zone, normalizing neuropeptide Y (NPY) release patterns during sympathetic excitation without altering norepinephrine (NE) release in the scar border zone. These findings extend the therapeutic window for cardiac afferent depletion beyond the acute phase of infarction^18,19^ and identify novel mechanisms by which afferent modulation confers antiarrhythmic benefit.

### Mechanistic insights

In this study, the antiarrhythmic effects of RTX were mediated through complementary structural, electrophysiologic, and neurochemical mechanisms. RTX-treated animals exhibited reduced border zone expansion and less conduction heterogeneity by electroanatomic mapping, consistent with the observed reduction in microstructural disarray and sympathetic nerve sprouting. High-resolution mapping revealed normalization of repolarization gradients and beat-to-beat ARI variability, further reflecting the stabilized post-infarct cardiac substrate. These findings indicate that not only is the scar border zone still being established 2-6 weeks after complete MI, but this period is also potently controlled by afferent neurotransmission. Given the expression of multiple cytokine receptors by nociceptive afferent nerve endings (e.g. receptors for TNF-α and IL1-β), persistent immune activation in the heart following MI perpetuates afferent neurotransmission. Our findings suggest that depleting afferent nerves following MI and by extension, eliminating the interactions between immune cells and nociceptive afferent nerves is cardioprotective.

An important mechanistic insight from this study is the demonstration that afferent depletion by RTX selectively restores evoked NPY release within the infarct border zone during sympathoexcitation, without affecting NE release. While it is known that persistent NE and NPY release is a feature of chronic ischemic cardiomyopathy,^30–32^ our finding of reduced NPY release in the scar-border zone of MI + Saline animals relative to sham and MI + RTX animals likely reflects this persistent release and depleted NPY stores, such that upon further sympathetic stimulation, less NPY is available for release. Conversely, in RTX-treated MI animals, the reduced persistent sympathetic drive (and consequently non-depleted NPY stores) result in significant NPY release on sympathetic stimulation. Interestingly however, afferent depletion modulates NPY but not NE release. While this finding may reflect differential remodeling of sympathetic neurotransmission after MI (i.e. NPY vs NE release), it may also reflect the reuptake mechanisms available for NE but not NPY, such that during states of chronic sympathetic excess, NE but not NPY remains available for evoked release. Independent of the mechanism, these findings reflect loss of adrenergic reserve and contribute to electrical instability, particularly in the highly arrhythmogenic border zone.

### Neuro-cardiac axis remodeling

It is recognized that following MI structural and functional alterations of intramyocardial nerves and the entire cardiac neuraxial control centers are central to neurohormonal activation, arrhythmias, and ventricular dysfunction.^5,33–36^ Our data demonstrate that cardiac-selective RTX treatment exerts effects beyond the myocardium, suppressing DRG neuroinflammation and downregulating immune response genes, thereby attenuating pro-excitatory afferent input. In the stellate ganglia, RTX reduced transcription of TH and NPY while upregulating pathways associated with axon guidance and extracellular matrix organization, suggesting a reprogramming of sympathetic neurons toward a less pro-arrhythmic phenotype.

These finding complement a recently described heart-brain-immune loop in which TRPV1-expressing vagal sensory neurons interact with angiotensin-II type-1 receptor neurons in paraventricular nucleus of the hypothalamus to modulate sympathetic tone in a mouse MI model.^14^. Ablating the sensory, central, or sympathetic node of this circuit limited infarct size, electrophysiologic abnormalities, and cardiac dysfunction. Another studying showed that epidural cervicothoracic RTX in a porcine chronic MI model restored vagal tone and baroreflex sensitivity, reduced spinal cord T cell infiltration and glial activation, stabilized electrophysiological parameters, and lowered VT/VF inducibility.^24^ Taken together, these findings highlight how potent cardiac nociceptive activation is following MI. These data indicate that persistent nociceptive afferent signaling after MI, whether relayed centrally through the vagus or spinal afferents, promotes coordinated remodeling of sensory and sympathetic neural circuits, initiating autonomic imbalance and supporting long-term electrical instability. RTX-mediated afferent depletion limited to the heart restores autonomic balance and electrical stability.

### Study limitations

Several limitations should be acknowledged. Although the porcine model closely recapitulates human cardiac anatomy and post-MI remodeling, validation in clinical cohorts will be required. While RTX selectively depleted TRPV1 afferents, potential long-term effects on pain pathways or other sensory modalities remain to be evaluated. While we demonstrate no significant loss of sensory neuron cell bodies in the DRG, the degree to which afferent signaling is restored following RTX treatment is unclear. Finally, although our data link restored NPY release and reduced arrhythmogenesis, direct causal relationships between NPY signaling and electrophysiologic stabilization remain to be mechanistically dissected.

### Conclusions

Cardiac-selective nociceptive afferent depletion with epicardial RTX, even when administered two weeks after a completed MI, prevents adverse remodeling, suppresses arrhythmias, and restores adrenergic reserve in the infarct border zone. These findings identify afferent modulation as a novel therapeutic strategy for post-MI arrhythmia risk reduction and support further translational development of targeted neuromodulatory therapies.

## CLINICAL PERSPECTIVES

### Core competencies

Previous studies have shown that RTX applied at the time of MI can attenuate adverse remodeling and reduce arrhythmia susceptibility.^18,19^ Our work extends these findings to the subacute post-MI period, when structural and electrophysiologic remodeling is already established, making this approach more clinically relevant. VT/VF remains a leading cause of mortality after MI despite current therapies. Standard antiarrhythmic agents are limited by systemic toxicity, while catheter ablation and device therapy primarily provide secondary prevention. Our findings position cardiac afferent depletion as a disease-modifying strategy that directly targets the arrhythmogenic substrate, stabilizes scar-border zone electrophysiology, and normalizes neurochemical signaling. Importantly, RTX preserved efferent autonomic reflexes, which may mitigate adverse hemodynamic consequences associated with global autonomic denervation.

### Translational outlook

This work addresses a critical therapeutic gap; a substantial proportion of patients with MI present after infarct completion, when opportunities for myocardial salvage are limited. By demonstrating that targeted afferent depletion restores adrenergic reserve specifically in the scar-border zone and reverses pro-arrhythmic remodeling, our data suggest that RTX therapy could be deployed during the early post-MI hospitalization or follow-up period to prevent arrhythmias and progression of heart failure. The percutaneous epicardial approach we applied here is technically feasible and compatible with existing interventional workflows, and the preservation of efferent sympathetic and parasympathetic function supports its safety profile. The ability to measure real-time neurotransmitter release *in vivo* also provides a framework for future biomarker development and patient selection in clinical translation.

## ACKNOWLEDGEMENTS

The authors thank Steven Cha, BS, Janaki Patel, BS, Sahil Haridas, BS, Nhi Doan, BS, Annie Kang, BS, and Kirellos Mikhail Atef, MD, for excellent technical assistance, and Kalyanam Shivkumar for comments on the manuscript.

## ABBREVIATIONS

ARI: activation recovery interval
AT: activation time
AWTd: anterior wall thickness at diastole
AWTs: anterior wall thickness at systole
CGRP: calcitonin gene-related peptide
CL: cycle length
CSAR: cardiac sympathetic afferent reflex
CsCl: cesium chloride
DRG: dorsal root ganglia
ECG: electrocardiogram
ERP: effective refractory period
GAP-43: growth associated protein 43
LVEF: left ventricular ejection fraction
LVIDd: left ventricular internal diameter at end-diastole
MI: myocardial infarction
NPY: neuropeptide Y
NSVT: non-sustained ventricular tachycardia
PES: program electrical stimulation
PGP 9.5: protein gene product 9.5
PVC: premature ventricle contraction
RSNS: right sympathetic nerve stimulation
RTX: resiniferatoxin
SD: standard deviation
SEM: standard error of the mean
SG: stellate ganglia
TH: tyrosine hydroxylase
TRPV1: transient receptor potential cation subfamily V member 1
VNS: vagal nerve stimulation

